# An *ex vivo* human precision-cut lung slice platform provides insight into SARS-CoV-2 pathogenesis and antiviral drug efficacy

**DOI:** 10.1101/2023.04.18.537373

**Authors:** Roger D. Pechous, Priyangi A. Malaviarachchi, Srijon K. Banerjee, Stephanie D. Byrum, Duah H. Alkam, Alireza Ghaffarieh, Richard C. Kurten, Joshua L. Kennedy, Xuming Zhang

**Author notes:** Corresponding author: Xuming Zhang.

## Abstract

COVID-19 has claimed millions of lives since the emergence of SARS-CoV-2, and lung disease appears the primary cause of the death in COVID-19 patients. However, the underlying mechanisms of COVID-19 pathogenesis remain elusive, and there is no existing model where the human disease can be faithfully recapitulated and conditions for the infection process can be experimentally controlled. Herein we report the establishment of an *ex vivo* human precision-cut lung slice (hPCLS) platform for studying SARS-CoV-2 pathogenicity and innate immune responses, and for evaluating the efficacy of antiviral drugs against SARS-CoV-2. We show that while SARS-CoV-2 continued to replicate during the course of infection of hPCLS, infectious virus production peaked within 2 days, and rapidly declined thereafter. Although most proinflammatory cytokines examined were induced by SARS-CoV-2 infection, the degree of induction and types of cytokines varied significantly among hPCLS from individual donors, reflecting the heterogeneity of human populations. In particular, two cytokines (IP-10 and IL-8) were highly and consistently induced, suggesting a role in the pathogenesis of COVID-19. Histopathological examination revealed focal cytopathic effects late in the infection. Transcriptomic and proteomic analyses identified molecular signatures and cellular pathways that are largely consistent with the progression of COVID-19 in patients. Furthermore, we show that homoharringtonine, a natural plant alkaloid derived from *Cephalotoxus fortunei*, not only inhibited virus replication but also production of pro-inflammatory cytokines, and ameliorated the histopathological changes of the lungs caused by SARS-CoV-2 infection, demonstrating the usefulness of the hPCLS platform for evaluating antiviral drugs.

**SIGNIFICANCE:** Here we established an *ex vivo* human precision-cut lung slice platform for assessing SARS-CoV-2 infection, viral replication kinetics, innate immune response, disease progression, and antiviral drugs. Using this platform, we identified early induction of specific cytokines, especially IP-10 and IL-8, as potential predictors for severe COVID-19, and uncovered a hitherto unrecognized phenomenon that while infectious virus disappears at late times of infection, viral RNA persists and lung histopathology commences. This finding may have important clinical implications for both acute and post-acute sequelae of COVID-19. This platform recapitulates some of the characteristics of lung disease observed in severe COVID-19 patients and is therefore a useful platform for understanding mechanisms of SARS-CoV-2 pathogenesis and for evaluating the efficacy of antiviral drugs.

## INTRODUCTION

The human respiratory tract is the primary target of infection by the severe acute respiratory syndrome coronavirus 2 (SARS-CoV-2) that causes the pandemic coronavirus disease 2019 (COVID-19). Clinical outcomes of SARS-CoV-2 infection vary widely from asymptomatic to death, and its underlying mechanisms remain elusive. The innate immune system in the respiratory tract is the first line of defense against invading respiratory pathogens, and infection often results in immediate and rapid induction of inflammatory cytokines, which in turn recruit leukocytes to infected sites that cause local inflammation and immunopathology. The speed, amount and type of local cytokines induced upon initial contact of a virus with target cells often dictates the outcome of infection. Indeed, clinical data have shown that the severity of COVID-19 often correlates with the onset of a pro-inflammatory cytokine storm and widespread alveolar damage and pneumonia (1), with the resulting lung disease the primary cause of death in patients (2, 3). However, because many social and behavioral factors influence SARS-CoV-2 transmission and disease outcome, what specific factors contribute to the initiation and progression of COVID-19 remains unclear. To better understand COVID-19 pathogenesis, it is critically important to have a model system where disease in humans can be recapitulated and the conditions for the infection process can be experimentally controlled.

Although animal models have been widely used for studying disease pathogenesis and preclinical testing for vaccines and therapeutics, there is always uncertainty as to what extent findings in animals recapitulate host–pathogen interactions in humans. Furthermore, the applicability of many animal models to human disease has been of concern, as countless findings have not translated during human clinical trials (4–8). While several animal models for SARS-CoV-1 and SARS-CoV-2 have been established, each model exhibits only certain clinical manifestations and histopathological features and does not faithfully reflect the whole picture observed in humans (9–11). Thus, developing an approach that can recapitulate SARS-CoV-2 infection in human lungs is critical for understanding its pathogenesis in humans and complementing the currently available infection models.

Though advances in tissue engineering have allowed for improved human infection models, advanced study of human lung tissue has been limited to the use of human primary and established cell lines (12). Since the emergence of the COVID-19 pandemic, several reports have described the development or adaptation of cell-or organoid-based systems, such as human stem cell-based alveolospheres and lung organoids for SARS-CoV-2 infection and antiviral drug screening (13–15). While these systems are permissive for SARS-CoV-2 infection, they lack the native lung environment and host cell repertoire. Analysis of primary human tissue has largely been limited to post-mortem analysis of samples from infected patients. We posit that engineering and fabrication of standardized platforms from viable human lungs obtained from deceased donors offer a critical native context for studying infectious diseases of the human lung. Human precision-cut lung slices (hPCLS) are slices of living human pulmonary tissue that can be maintained under standard cell culture conditions in a laboratory. The hPCLS platform maintains the 3D cellular structure present in native tissue, and therefore fills a critical gap in existing infection models. The hPCLS platform accurately reflects not only the actual lung niche, preserving ciliary beat frequency and mucous production, but also cellular viability of the entire repertoire of cells found in the lung, including alveolar epithelial cells, endothelial cells, dendritic cells, alveolar and interstitial macrophages, and type 2 innate lymphoid cells (16). It also elicits diverse cytokine and chemokine responses and airway hyperresponsiveness to infection (17, 18). The complexity of human lung tissue supports direct translation of results from animal to human and from *in vitro* to *in vivo*. Herein we report the establishment of the *ex vivo* hPCLS platform as a powerful tool for studying SARS-CoV-2 pathogenicity and innate immune response, and for evaluating the efficacy of antiviral drugs against SARS-CoV-2.

## RESULTS

### Establishment of the hPCLS platform for SARS-CoV-2 infection

As human lungs are the native organ for SARS-CoV-2 infection, we sought to establish the *ex vivo* hPCLS platform for studying SARS-CoV-2 infection. Transplant-quality lungs (**Fig. 1A-a**) were processed into hPCLS that can be maintained in the laboratory for up to 3 months (**Fig. 1A-b,c**). hPCLS slices were infected with SARS-CoV-2 and supernatants were collected at 3 and 24 h p.i. for determining virus titer. We found that SARS-CoV-2 indeed replicated in hPCLS as viral titers increased to 2×10^4^ TCID_50_/ml from 3 to 24 h p.i. (**Fig. 1B**). We then determined viral RNAs in the hPCLS using qRT-PCR. Consistent with the virus titers, viral RNAs were increased >360 fold over mock-infected control at 24 h p.i. (**Fig. 1C**). Viral N protein was also detected in hPCLS at 24 h p.i. by immunofluorescence (**Fig. 1D**). It is noted that the immunofluorescence staining was generally weak and the majority of the N protein appeared in type II alveolar cells (**Fig. 1D**).

**Fig. 1.**
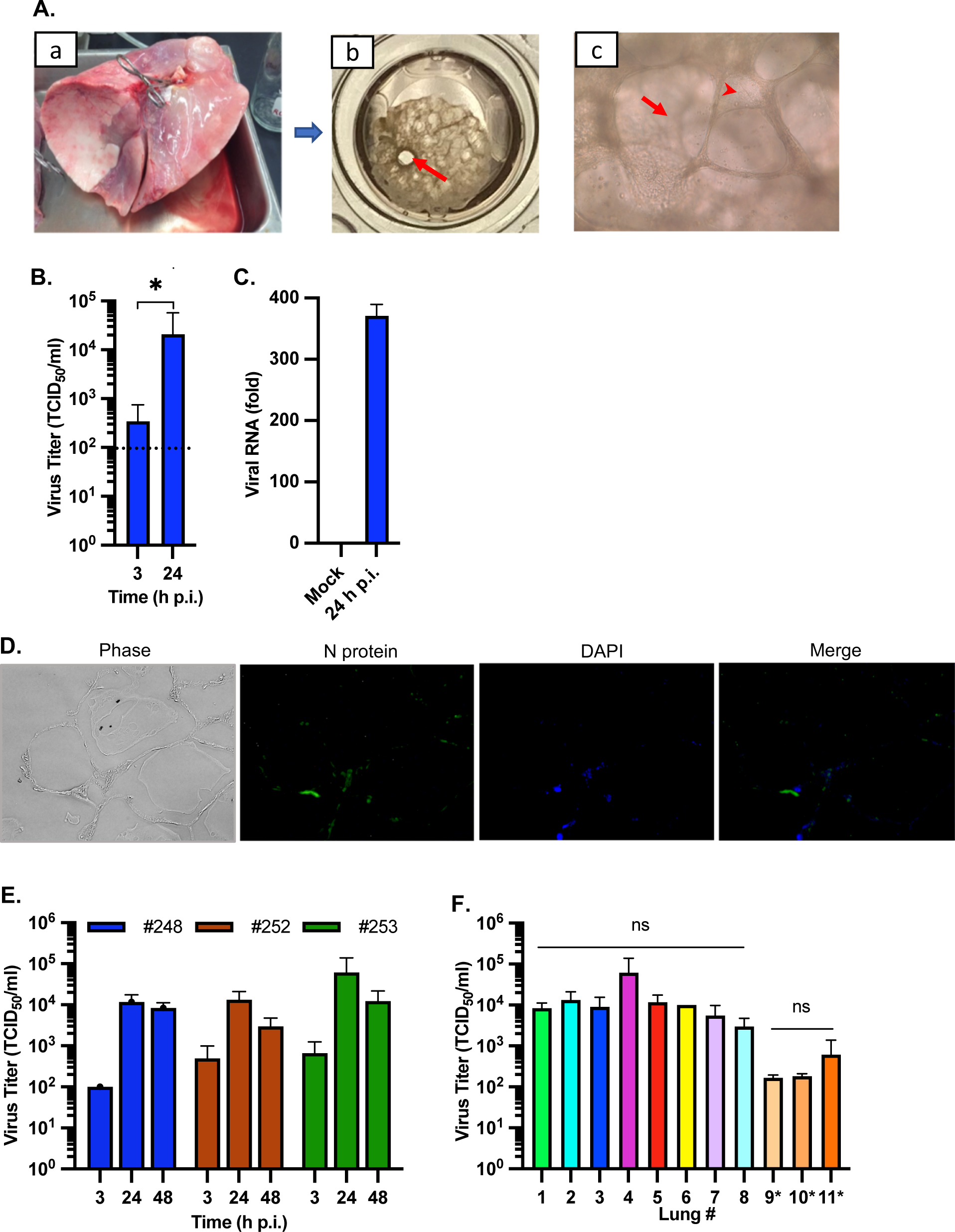
Establishment of the hPCLS platform for SARS-CoV-2 infection. (A) Processing of hPCLS. Donated human lungs (a) were processed into hPCLS (b, c). (b) An image of hPCLS with one or more large airways (arrow) in one well of a 48-well plate. (c) Bright field image of hPCLS at 200x magnification showing alveoli (arrow) and interstitial spaces (arrowhead). (B) Analysis of susceptibility of hPCLS to SARS-Cov-2 infection. hPCLS were infected with SARS-CoV-2 at 2×10^5^ TCID_50_ per slice, and culture supernatants were collected for determination of virus titer (in TCID_50_/ml) at 3 and 24 h p.i. Data is representative of 3 independent experiments and is the mean and standard deviation (SD) of a triplicate. *, *P*<0.05 (unpaired T test). (C) Quantification of viral RNA (in fold) in hPCLS at 24 h p.i. relative to mock-infected control as measured by qRT-PCR and expressed as the mean and SD of a duplicate. (D) Detection of SARS-COV-2 N protein by immunofluorescence. hPCLS were infected with SARS-CoV-2 at 2×10^5^ TCID_50_ per slice for 24 h. Slices were fixed with formalin and processed and embedded with paraffin. The slices were then stained with a monoclonal antibody against viral N protein and an anti-mouse IgG conjugated with FITC. Cell nuclei were stained with DAPI. (E-F) Susceptibility of hPCLS derived from different individual. hPCLS were infected with SARS-CoV-2 for a period of time as indicated (E) or for 24 h (lung#1-8) or 48 h (lung#9-11 with asterisk) (F). The culture supernatants were harvested for determination of virus titer. Data is the mean and SD of 3 replicates for each individual lung as indicated. ns, *P*>0.05 (one way ANOVA).

To determine if individuals have different susceptibilities to SARS-CoV-2 infection, hPCLS from 3 donors were infected with SARS-CoV-2 for 3, 24 and 48 h. Results show that the difference in virus titers between the 3 donor lungs was relatively small, i.e., within 1 log_10_ (**Fig. 1E**). Virus titers were also similar in 8 additional donors at 24 h p.i. or 3 donors at 48 h p.i. (**Fig. 1F**). It is noted that the ages of these donors range from 31 to 48 years. While the number of donors is small, these results suggest that lungs from adults between 30 and 50 years of age have similar susceptibility to SARS-CoV-2 infection. Collectively, these results demonstrate that the *ex vivo* hPCLS platform is permissive for SARS-CoV-2 infection.

### Kinetics of SARS-CoV-2 replication in hPCLS

Little is known about the precise viral replication kinetics in the lungs of individual COVID-19 patients. This information is critical for understanding COVID-19 pathogenesis and for effectively managing COVID-19 patients. We took advantage of the hPCLS platform to determine the replication characteristics of SARS-CoV-2 in human lungs under controlled experimental conditions. hPCLS slices were infected with SARS-CoV-2 and culture supernatants were collected for determining virus titers. As shown in Fig. 2A, virus titers rapidly increased from 3 to 24 h p.i., reached a plateau at 1.2×10^4^ TCID_50_/ml at 36 h p.i., and rapidly decreased thereafter. By 96 h p.i. infectious viruses were barely detectable (167 TCID_50_/ml), suggesting transient reproduction kinetics for infectious viruses. However, viral RNAs in the lungs increased continuously from 24 to 96 h p.i. (**Fig. 2B**). This result is in stark contrast to those obtained from Vero and human lung epithelial A549/ACE2 cell culture systems, in which virus replication reached and maintained high titers (≈10^7^ TCID_50_/ml) from 24 to 72 h p.i., and decreased only slightly (≈1 log_10_) from 72 to 96 h p.i. (**Fig. 2C**). These results indicate that virus titers increased rapidly in both immortalized cell lines and hPCLS during the first 24 h of infection, followed by a plateau, and in the case of hPCLS decreased rapidly.

**Fig. 2.**
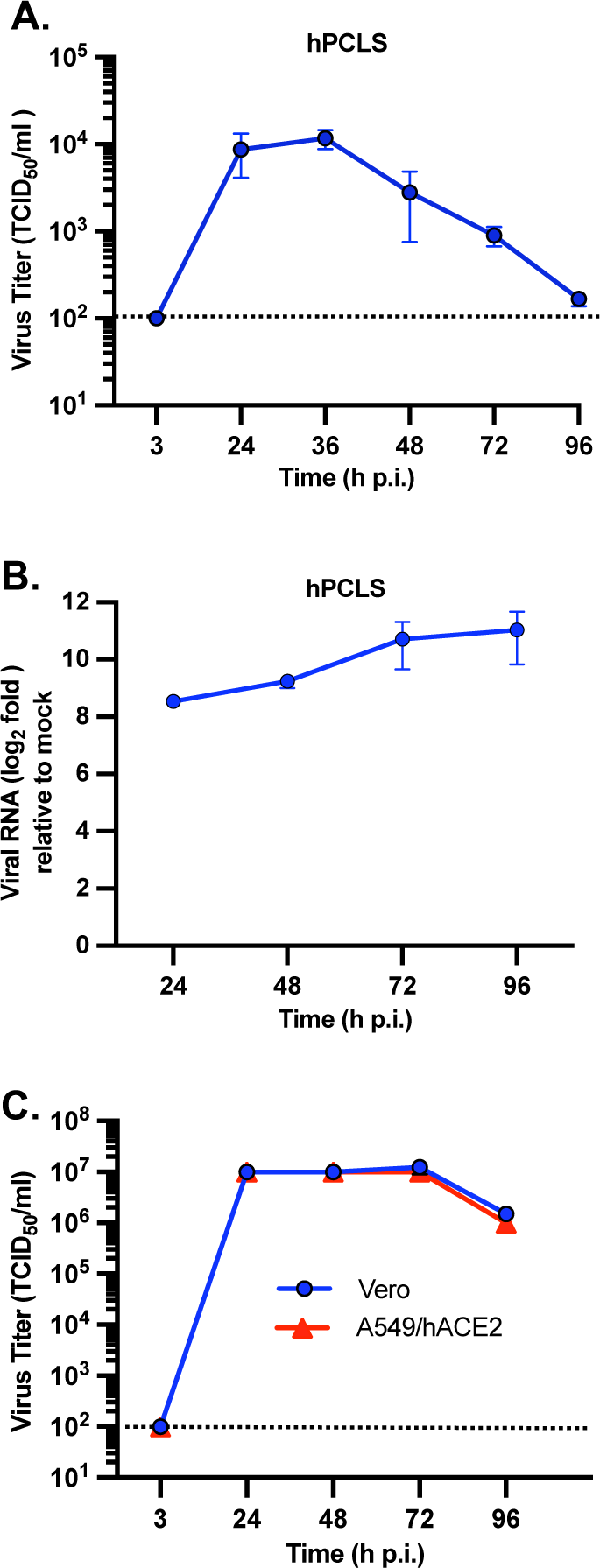
SARS-CoV-2 replication kinetics in hPCLS. hPCLS were infected with SARS-CoV-2 at2×10^5^ TCID_50_ per slice. At indicated time p.i., supernatants were harvested for determining virus titer (A) and slices for viral RNA quantification (B). Virus titer is expressed in TCID_50_/ml and viral RNA in log2 fold over the mock-infected control. Data is the mean and standard deviation (SD) of 3-6 replicates. (C) SARS-CoV-2 replication kinetics in cell cultures. Vero or A549/ACE2 cells were infected with SARS-CoV-2 at MOI of 1 and supernatants were harvested at various time points p.i. for determination of virus titer, which is expressed as the mean TCID_50_/ml of a duplicate and SD of the means.

### Induction of proinflammatory cytokines and chemokines in hPCLS by SARS-CoV-2 infection

The innate immune response in the respiratory tract is the first line of host defense against respiratory pathogens, but it can also trigger damaging inflammation. Initial induction of local cytokines and chemokines often dictate the outcome of an infection or disease. To identify the initial innate immune response to SARS-CoV-2 infection in the lungs that might drive progression of COVID-19, we assessed the induction of common proinflammatory cytokines and chemokines in hPCLS following SARS-CoV-2 infection. We identified several features of the proinflammatory response in the lungs (**Fig. 3**). First, three cytokines and chemokines (IL-8, IP-10, and MCP-1) were highly induced (ranging from 946 to 7,350 pg/ml at 48 h p.i.) while four cytokines and chemokines (IL-1β, TNF-α, RANTES, MIG) were only modestly induced (ranging from 2 to 69 pg/ml at 48 h p.i.) by SARS-CoV-2 infection. Second, there was generally a continuous increase in expression during the first 48 h of infection. Third, there was a significant variation in both specific cytokines and the level of induction between individual donor lungs. For example, MCP-1 and TNF-α were not induced in donor #248 and donor #252, respectively, while RANTES was not induced in both donors #248 and #253. IL-8 was induced to 7,350 pg/ml in donor #248 but only to 1,393 pg/ml in donor #252 at 48 h p.i. Fourth, four cytokines and chemokines (IL-8, IP-10, MIG, IL-1β) were induced in all three donor lungs, with IP-10 and IL-8 being at high levels. Fifth, IL-10 and IL-12p70 were not induced by SARS-CoV-2 infection; however, induction of IL-6 appeared nonspecific as it was induced in both virus-infected and mock-infected hPCLS (data not shown). We thus conclude that IP-10 and IL-8 are two reliable inflammatory biomarkers for SARS-CoV-2 infection in the lungs as both were consistently induced in all donors at high levels.

**Fig. 3.**
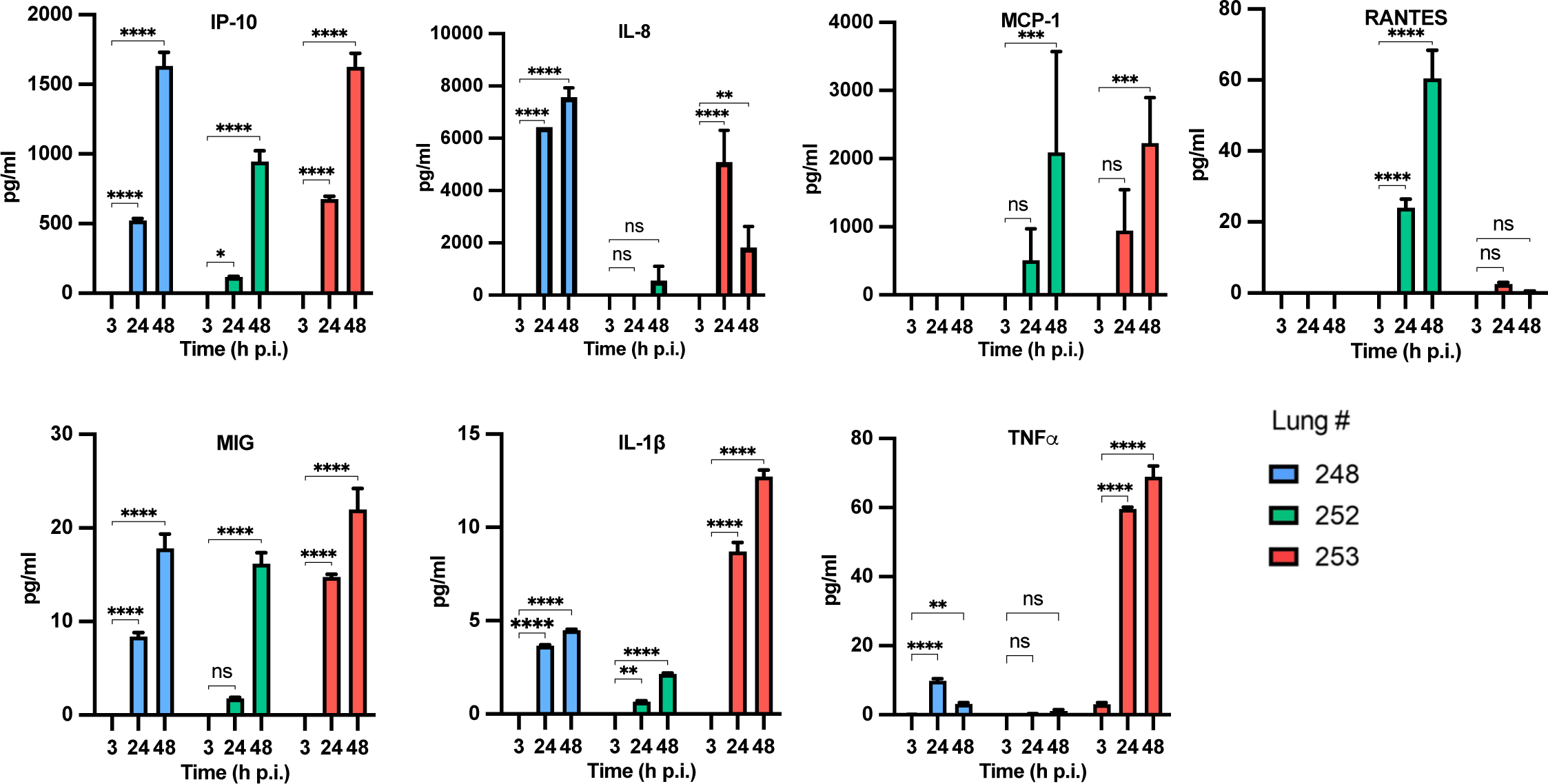
Induction of proinflammatory cytokines and chemokines in hPCLS by SARS-CoV-2 infection. hPCLS were infected with SARS-CoV-2 at 2×10^5^ TCID_50_ per slice, and culture supernatants were harvested at 3, 24, and 48 h p.i. for quantifying the protein levels of secreted cytokines and chemokines using cytometric beads array (CBA) kits by flow cytometry. The amounts from infected samples were subtracted by the amounts from mock-infected samples, and were expressed as pg/ml. Data is the mean and SD of 3 replicates for each cytokine and chemokine and is indicated for 3 individual lung donors (#248, #252, #253). Significance was calculated with Tukey’s multiple comparisons test in the GraphPad Prism program. *, *P* < 0.05; **, *P* < 0.01; ***, *P* < 0.001; ****, *P* < 0.0001.

### Development of histopathology in hPCLS following SARS-CoV-2 infection

Histopathological changes in the lungs can result from direct virus infection (i.e., cytolytic infection) or a bystander effect (i.e., through immune response). Induction of proinflammatory cytokines often leads to inflammation that in turn results in histopathological changes in the lungs, which is a hallmark of severe COVID-19 in patients (19). To evaluate the utility of the hPCLS platform for studying COVID-19 pathogenesis, we determined histopathological changes of hPCLS following SARS-CoV-2 infection. As shown in Fig. 4, despite the induction of proinflammatory cytokines as early as 24 h p.i. and a significant decrease in viral titers at late times (48-96 h) p.i., localized (focal) cytopathic effects as indicated by cellular debris in alveolar spaces weren’t observed in SARS-CoV-2-infected hPCLS until day 5 p.i. (marked areas). These results suggest that the onset of a proinflammatory response results in a cytopathic effect in host tissue, albeit delayed until roughly 5 d p.i.

**Fig. 4.**
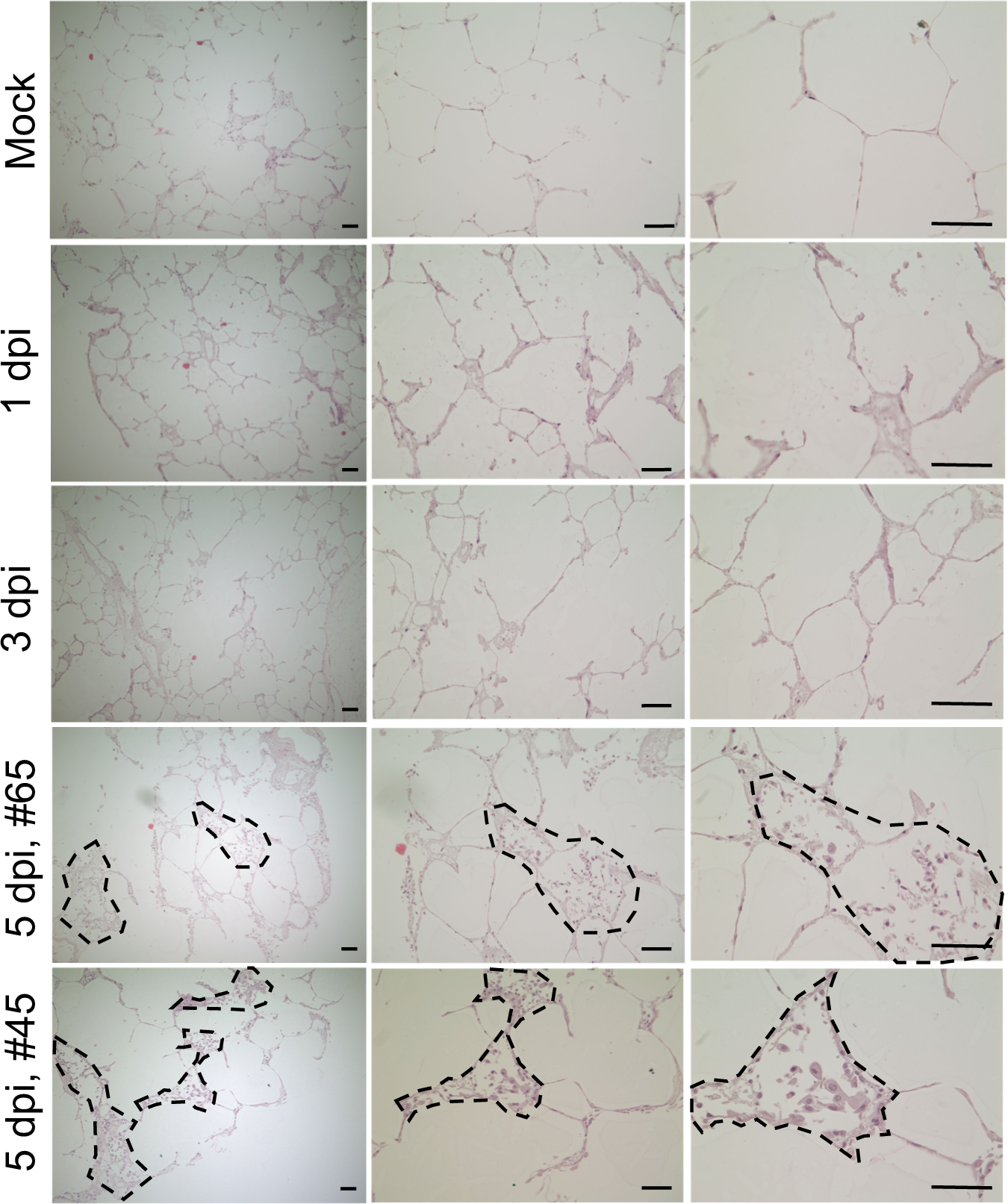
Histopathological development in SARS-CoV-2-infected hPCLS. hPCLS were infected with SARS-CoV-2 at 2×10^5^ TCID_50_ per slice for 1, 3 and 5 days or mock-infected as a control, and the slices were fixed and processed for H&E staining. Areas with cytopathic effects at day 5 p.i. are marked with dashed lines in 2 hPCLS samples (#45 and #65). Data are representative of the experiments from 6 individual lungs. Scale bar, 50 µm.

### Gene signatures identified by transcriptomic profiling shed light on COVID-19 progression in the lungs

While gene expression profiles from the lungs of post-mortem COVID-19 patients have been reported (20), these data represent only a snap shot of the disease and do not reflect the disease progression in individual patients during infection. To assess early host responses in the lung that drive COVID-19 progression, we determined the transcriptome profiles in hPCLS from 24 to 96 h following infection by RNAseq. We performed differential and pathways analyses and found that at 24 h p.i. 87 genes were upregulated while 433 genes were downregulated by SARS-CoV-2 infection with a p-value <0.05 and a log2 fold change >1 (**Fig. 5A**). The number of genes that were up-regulated at the 48 h, 72 h and 96 h p.i. were 358, 744, and 822, respectively. The number of genes that were down-regulated at the 48 h, 72 h and 96 h p.i. were 237, 995, and 2233, respectively (**Fig. S1 and Dataset S1-S4**). Of note, three of the top 10 down-regulated genes were *TNFSF11*, *CA4* and *OSGIN1*, which are involved in T cell-dependent immune response, CO_2_/O_2_ exchange, and NRF2-dependent antioxidant gene expression and cell death, respectively. Down-regulation of these genes may repress T cell immunity, decrease lung function, and exacerbate oxidative stress and cell death. On the other hand, up-regulation of chemokine CCL4/MIP-1β, PPFIA4, and SFTPA2 (surfactant protein A2) that are involved in regulation of inflammation, cell adhesion, and interstitial lung disease/pulmonary fibrosis, respectively, potentially promotes inflammation and lung disease. Five KEGG pathways were particularly enriched (**Fig. 5B**), indicating commencement of an inflammatory phase. By 48 h p.i., while more genes involved in cell adhesion and migration were continuously and significantly up-regulated, enriched genes in two additional pathways (i.e., HIF-1 and cellular senescence) began to emerge (**Fig. 5C**), suggesting that hypoxia and senescence have initiated at this stage. However, many of the genes in IL-17 signaling pathways and viral protein interaction with cytokine and cytokine receptor pathway were downregulated (**Fig. 5D**). In particular, *CCL17* was downregulated more than 7 log2 folds. These results indicate a transient nature of transcriptional regulation of these chemokine genes in infected lungs. At 72 h p.i., a large cluster of 35 genes related to ribosome were upregulated, all of which have been previously implicated in COVID-19 (**Fig. 5E**). Additionally, a cluster of 25 enriched genes in the protein processing in endoplasmic reticulum pathways were significantly upregulated, including many of the heat shock proteins. Twelve genes enriched in the p53 signaling pathway were also upregulated (**Fig. 5E**). By 96 h p.i., enriched genes in 3 pathways were upregulated, including 13 genes in mitophagy, 10 genes in ferroptosis and 14 genes in complement and coagulation cascades (**Fig. 5F**). Of particular note are *MAP1LC3C* (>6 log2 fold increase) and *SERPINA5* (>6 log2 fold increase) that are involved in cell death and blood coagulation, respectively. These results indicate that SARS-CoV-2 infection leads to profound changes of the transcriptional landscape in the lungs early during infection.

**Fig. 5.**
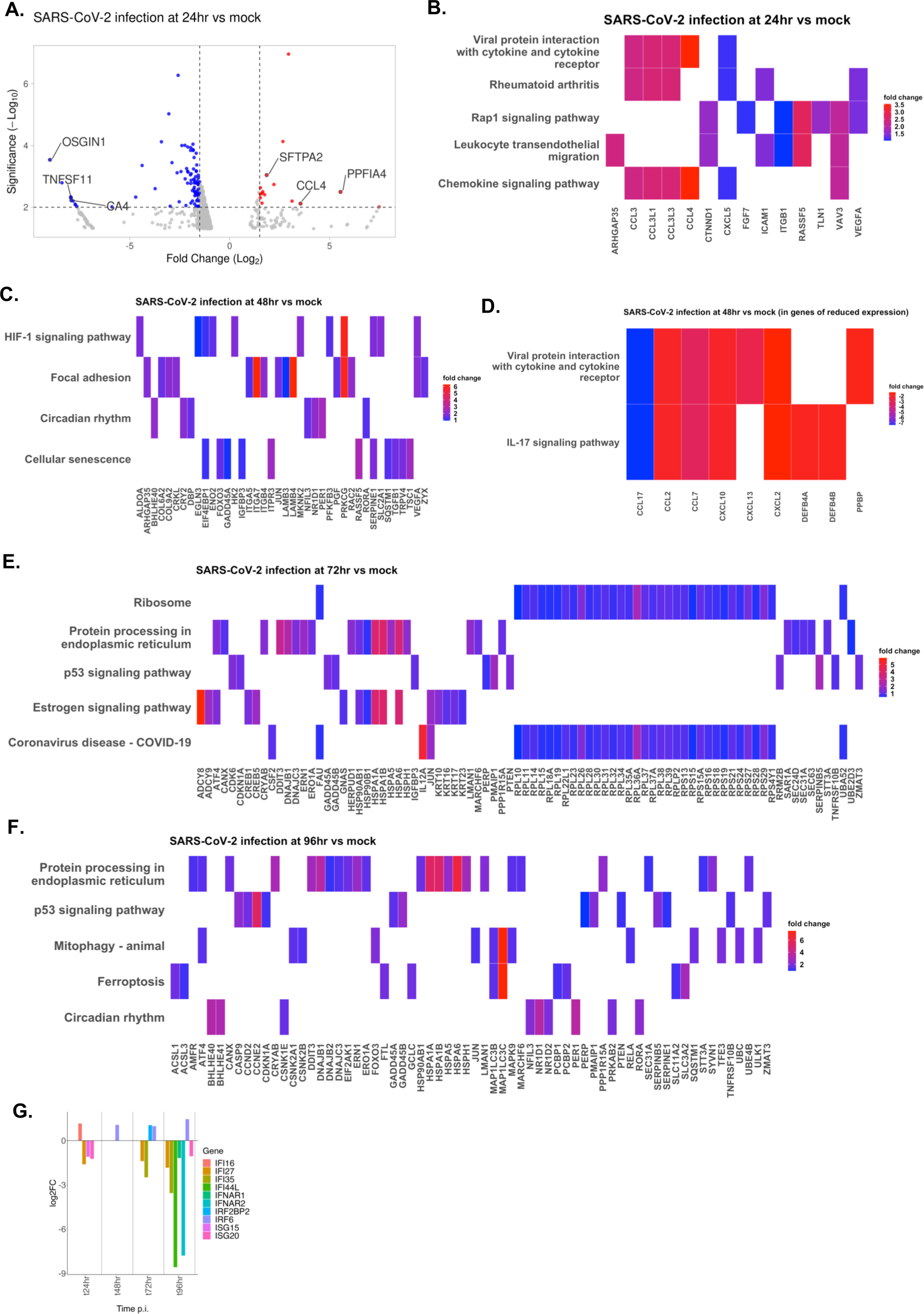
Gene signatures in COVID-19 progression in the lungs identified by transcriptome profiling. Groups of 6 hPCLS slices each were infected with SARS-CoV-2 at 2×10^5^ TCID_50_ per slice or mock-infected as a control. At various time points (24-96 h) p.i., RNAs were isolated from the slices and subjected to RNAseq analysis. Differential gene expression was identified by comparing virus-infected group to mock-infected group. (A) A volcano plot showing the down-and up-regulated genes in the lungs by SARS-CoV-2 at 24 h p.i. with 3 representative signature genes each. (B)-(F) Heat maps showing signature genes in the major pathways during the disease progression in the lungs at 24 (B), 48 (C and D), 72 (E) and 96 (F) h p.i. (G) Bar plot showing the differential expression of IFN-I- and IFN-II-related genes at 24, 48, 72 and 96 h p.i. Positive and negative log2FC values indicate upregulation and downregulation, respectively.

Notably, 7 type I interferon (IFN-I) related genes were significantly down-regulated (up to −8 log2 fold reduction) at various time points p.i., and none of the other IFN-I related genes were up-regulated at any of the 4 time points p.i. (**Fig. 5G, Dataset S1-S4**). In contrast, 2 type II interferon (IFN-II)-related genes (*IRF6, IFI16*) were moderately upregulated (up to 1.4 log2 fold increase). *IRF2BP2*, which is an interferon regulatory factor 2 (IRF2)-binding protein and acts as a co-repressor for IRF2 to repress IFN-I gene transcription (21), was also moderately up-regulated (1 log2 fold increase) at 72 h p.i. (**Fig. 5G**).

### Proteomic analysis provides insight into activation of host molecular networks during SARS-CoV-2 infection in human lungs

To further understand the molecular basis of SARS-CoV-2 pathogenesis in human lungs, we determined the proteomic profiles in hPCLS at 48 h p.i., and identified a number of upstream molecules, regulators, pathways and diseases that are associated with SARS-CoV-2 infection (**Fig. 6A**). Specifically, several growth factors, cytokines and chemokines networks appeared to be activated, which leads to branching of vasculature, infiltration by T lymphocytes, and cell movement of leukocytes, all indicative of the onset of an inflammatory phase of the disease, consistent with the findings from RNAseq (**Fig. 5**). Activation of these growth factors and many other transcriptional regulators (e.g., MRTFA, MRTFB, FOXM1, NPM1 and TEAD4) can also lead to formation of intercellular junctions, sprouting, and inhibition of organismal death. These functional changes are predicted to trigger several signaling pathways, e.g., (hepatic) fibrosis, wound healing, and dilated cardiomyopathy. Notably, the canonical GP6 signaling pathway is predicted to be activated by SARS-CoV-2 infection (**Fig. 6A**). GP6 is a member of the immunoglobulin superfamily and serves as the major signaling receptor for collagens and laminins, which leads to the platelet activation and thrombus formation. Indeed, numerous collagens and laminins were significantly activated in SARS-CoV-2-infected lungs (a partial list shown in **Fig. 6B**). The IPA also predicts potential links to several diseases, such as systemic lupus erythematosus, human papillomavirus infection, and alcoholism (**Fig. 6B**). These results suggest that the cellular proteomic networks in the lungs that are altered by SARS-CoV-2 during the first 48 h of infection not only promote pulmonary inflammation, but may also contribute to other aspects of COVID-19, such as fibrosis, heart failure, thrombosis, and autoimmune disease.

**Fig. 6.**
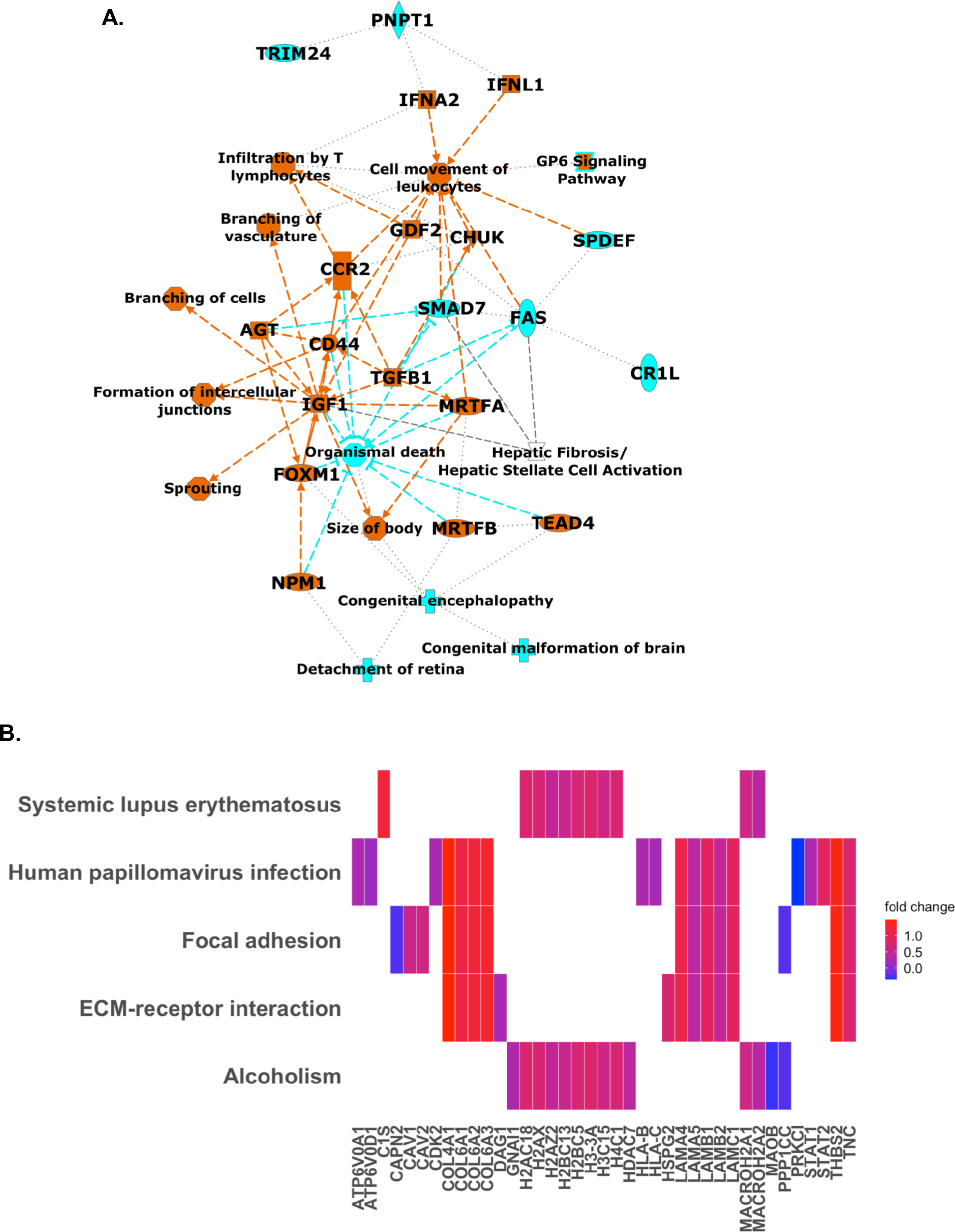
Alterations of molecular networks in human lungs during SARS-CoV-2 infection as determined by proteomic analysis. Groups of 6 hPCLS slices each were infected with SARS-CoV-2 at 2×10^5^ TCID_50_ per slice or mock-infected as a control. At 48 h p.i., proteins were isolated from the slices and subjected to proteomic analysis. The proteomes in virus-infected hPCLS were compared to those in mock-infected hPCLS. (A) Summary of upstream molecules, regulators, pathways and diseases that are associated with SARS-CoV-2 infection by Ingenuity Pathway Analysis (IPA). Node symbol: 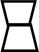 canonical pathway; 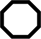 Function; cytokine; 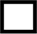 growth factor; 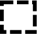 transcription regulator; 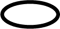 transmembrane receptor; 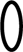 complex; 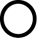 G-protein coupled receptor; 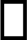 enzyme; 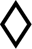 kinase; 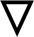 disease. 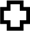 Color code: Orange, upregulation; blue, downregulation. (B) Major pathways and associated proteins in the lungs that are up-regulated by SARS-CoV-2 infection.

### The hPCLS platform as a “clinical trial at the bench” for evaluating antiviral drugs against SARS-CoV-2

Cell cultures and small animal models have been the gold standard for pre-clinical testing of antivirals. However, information gained from cell cultures are limited to antiviral effect and cytotoxicity of a drug. While animals are essential for *in vivo* testing, countless findings have not translated during human clinical trials (4–8). To extend the utility of the hPCLS platform, we carried out antiviral drug testing in hPCLS (**Fig. 7A**). We used homoharringtonine (HHT) as the first example. We previously identified HHT as a potent anti-coronavirus small molecule compounds during library screening (22), and recent studies have confirmed its anti-SARS-CoV-2 activity in cell culture (23) (**Fig. S2**). Our results from three individual donors confirmed that HHT at 1 µM had an antiviral activity against SARS-CoV-2 with a reduction of virus titer ranging from 2-to 14-fold (**Fig. 7B**). We also determined the antiviral effect of HHT at concentrations ranging from 1 to 10 µM. The results show that HHT completely inhibited SARS-CoV-2 replication starting at 5 µM (**Fig. 7C**).

**Fig. 7.**
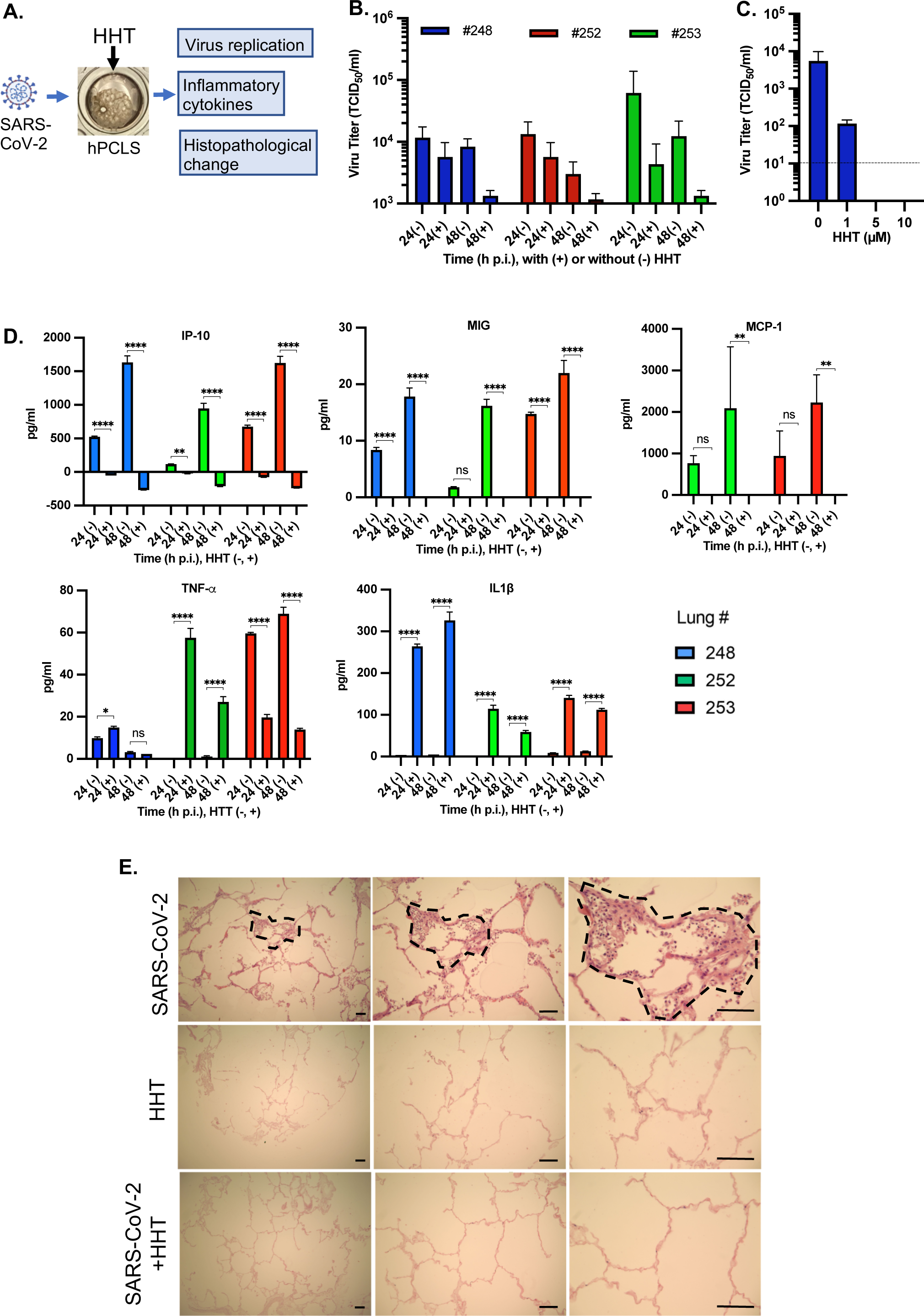
Establishment of the *ex vivo* hPCLS platform for evaluating antiviral drugs. (A) Schematic of the experimental design. HHT, homoharringtonine. (B, C) Inhibition of SARS-CoV-2 replication in hPCLS by HHT. (B) hPCLS from 3 individual lung donors as indicated were infected with SARS-CoV-2 at 2×10^5^ TCID_50_ per slice and treated with HHT at 1 h p.i. at 1 µM for 24 h (24+) or 48 h (48+) or untreated for 24 h (24-) or 48 h (48-). Virus titers in the supernatants were determined and expressed as TCID_50_/ml. Data is the mean and standard deviation (SD) of 3 replicates. (C) Determination of dose response. Six hPCLS each were infected with SARS-CoV-2 at 2×10^5^ TCID_50_ per slice or mock-infected, and treated with HHT at 1 h p.i. at various concentrations as indicated. Virus titers in the supernatants were determined and expressed as TCID_50_/ml. Data is the mean and SD of 3 replicates. (D) Inhibition of proinflammatory cytokines and chemokines by HHT. hPCLS were infected with SARS-CoV-2 at 2×10^5^ TCID_50_ per slice and treated with HHT (+) at 1 µM or untreated (-). Supernatants were harvested at 24 h or 48 h p.i., and the amounts of cytokines were determined with cytometric beads array assay. The amounts from infected samples were subtracted by the amounts from mock-infected samples, and were expressed as pg/ml. Data is the mean and SD of 3 replicates for each cytokine and chemokine and is indicated for 3 individual lung donors (#248, #252, #253). Significance was calculated with Tukey’s multiple comparisons test in the GraphPad Prism program. *, *P* < 0.05; **, *P* < 0.01; ***, *P* < 0.001; ****, *P* < 0.0001. (E) Inhibition of SARS-CoV-2-caused histopathological abnormality in hPCLS by HHT. hPCLS were mock-infected or infected with SARS-CoV-2 at 2×10^5^ TCID_50_ per slice and treated with HHT at 1 h p.i. at 10 µM. On day 5 p.i., slices were fixed with formalin and processed for H&E staining. Areas with cytopathic effects are marked with dashed lines. Data are representative of the experiments from 3 individual lungs. Scale bar, 50 µm.

We then determined the impact of HHT treatment on the induction of proinflammatory cytokines mediated by SARS-CoV-2 infection. Results show that induction of chemokines IP-10, MIG, and MCP-1 was completely blocked following treatment with HHT at 1 µM (**Fig. 7D**). TNF-α was not affected in donor #248, increased in donor #252, but decreased in donor #253 by treatment with HHT, indicating a significant variability among different donor lungs. In contrast, IL-1β was induced for all donors by HHT (**Fig. 7D**). The effect of HHT on the expression of IL-8 and RANTES was inconclusive (data not shown). Taken together, these results demonstrate that HHT has a general anti-inflammatory effect but that its effect on specific cytokines can vary between the individuals.

We further evaluated the impact of HHT treatment on histopathological changes of the lungs. hPCLS were infected with SARS-CoV-2 and treated with HHT at 10 µM for 5 days. As shown in Fig. 7E, localized cytopathic effects were observed in SARS-CoV-2-infected hPCLS (marked areas). In contrast, no cytopathic effect was observed in SARS-CoV-2-infected and HHT-treated hPCLS, as in mock-infected and HHT-treated hPCLS (**Fig. 7E**). These results indicate that treatment with HHT reversed the lung histopathology caused by SARS-CoV-2 infection, and that the induction of IL-1β by HHT likely did not contribute to the histopathological abnormality.

## DISCUSSION

In the present study, we established an *ex vivo* hPCLS platform for evaluating the pathogenesis of SARS-CoV-2 infection in human lungs. Although SARS-CoV-2, especially the omicron variant, is highly transmissible, a large proportion of SARS-CoV-2 infections result in mild or no clinical symptoms. The most severe and fatal cases of COVID-19 are primarily caused by lung diseases; yet how SARS-CoV-2 replicates in the lungs is not well understood. Although airway epithelial cells are the initial targets of SARS-CoV-2 infection, many cell lines derived from airway epithelium, e.g., A549 (type II alveolar cells) and BEAS-2 (bronchial epithelial cells), are not or are minimally susceptible to SARS-CoV-2 infection. Primary epithelial cells isolated from different regions of the respiratory tract have different levels of ACE2 (angiotensin-converting enzyme-2) receptor, which correlate with their susceptibility to SARS-CoV-2 infection (18, 24, 25). Thus, most cell culture models may not effectively recapitulate the characteristics of viral replication in the lungs (26). We show that SARS-CoV-2 infectivity in hPCLS as measured by virus titer (**Fig. 1**) differs from that seen in isolated primary cells, i.e., lower than in nasal or large airway epithelial cells but higher than in types I and II alveolar cells (18), indicating that the hPCLS platform represents mixed populations of different cell types in the native lungs.

Cell heterogeneity is a salient feature of the hPCLS platform. The hPCLS platform reflects not only the actual lung niche, preserving ciliary beat frequency and host cell responsiveness, but also cellular heterogeneity. As leukocytes play critical roles in defense against respiratory pathogens as well as in mediating lung inflammation, the presence of leukocytes in hPCLS provides an excellent *ex vivo* infection relevant to infection of actual human lungs. Indeed, a third of the cell populations in a typical hPCLS are CD45+ leukocytes/lymphocytes, with alveolar macrophages, dendritic cells, classical monocytes, and interstitial macrophages representing the bulk of the leukocyte populations (27). Furthermore, the hPCLS platform preserves the actual 3D architecture of the lungs, including the blood vessels and interstitial spaces (**Fig. 1A**). Because interactions and communications among different cell types are essential for maintaining lung homeostasis and in responses to pathogen invasion, the hPCLS platform offers a unique perspective for evaluating the pathogenesis of COVID-19 over recently reported *in vitro* models such as alveolospheres and lung organoids that lack the relevant cell heterogeneity or other components (13, 15). Supplementing additional peripheral blood mononuclear cells would make the hPCLS platform even more closely resemble the lungs *in vivo*.

We observed that the kinetics of infectious SARS-CoV-2 production in hPCLS does not correlate with that of viral RNA (**Fig. 2**). This is striking, because in general virus titer correlates with viral RNA accumulation during acute infection in a given host. Whether this phenomenon is unique to SARS-CoV-2 or to the hPCLS platform remains unclear. This finding raises several interesting questions. For example, do viral RNAs continue to replicate in the lungs without apparent production of infectious virus after 4 days? If so, how long does the viral RNA persist in the lungs and what are the consequences to the host? SARS-CoV-2 RNAs have been detected in patients long after recovery from acute infection (28–30) and in multiple postmortem organs including the brain as late as 230 days after onset of symptoms (31). Viral RNA persistence in the absence of infectious virus has also been described in oligodendrocytes and in mouse brains infected with murine coronavirus (32–34). The observation that cytopathic effects are found in hPCLS on day 5 p.i., when infectious virus could no longer be detected, suggests that continuous viral gene expression in the absence of infectious virus production can still result in histopathological changes of the lungs (**Fig. 4**). This finding may have important clinical implications in post COVID-19 sequelae and warrants further investigation.

While the precise mechanism by which SARS-CoV-2 causes lung disease is unknown, our analysis of this experimental infection platform paints an emerging picture of SARS-CoV-2 pathogenesis that can be divided into three phases (**Fig. 8**). During the initial phase of infection (first 48 h), SARS-CoV-2 replicates rapidly (**Fig. 2**) and triggers immediate innate immune responses, as evidenced by rapid induction of pro-inflammatory cytokines and chemokines and high levels of secreted IL-8 and IP-10 (**Figs. 3 & 8A**). Rapid induction of these local inflammatory cytokines likely plays a role in the development of lung disease, as supported by clinical evidence that shows that high levels of IL-8 and IP-10 in patients sera correlate with severity of COVID-19 (35–38). Thus, both local (lung) and peripheral IL-8 and IP-10 can be considered reliable predictors for progressive and severe COVID-19. Changes in hPCLS cellular gene expression in response to infection appear to promote cell growth, survival, and trafficking in the lung environment (**Figs. 5, 6 & 8B**), which is also characteristic of an early stage of inflammation. At the second phase (48-96 h), while infectious virus production rapidly decreases (**Fig. 2A**), viral RNA continues to accumulate (**Fig. 2B**). This may suggest that though infectious virus is cleared, continuous viral gene expression may have lingering effects on the host. Indeed, clusters of most enriched genes are involved in cellular pathways regulating cell death, such as the p53 signaling pathway, mitophagy, and ferroptosis (**Figs. 5F**), which may play a role in the development of histopathological abnormality at the late stage (third phase) of infection (5 days p.i.) (**Figs. 4 & 8**), and as observed in post-mortem COVID-19 patients (19).

**Fig. 8.**
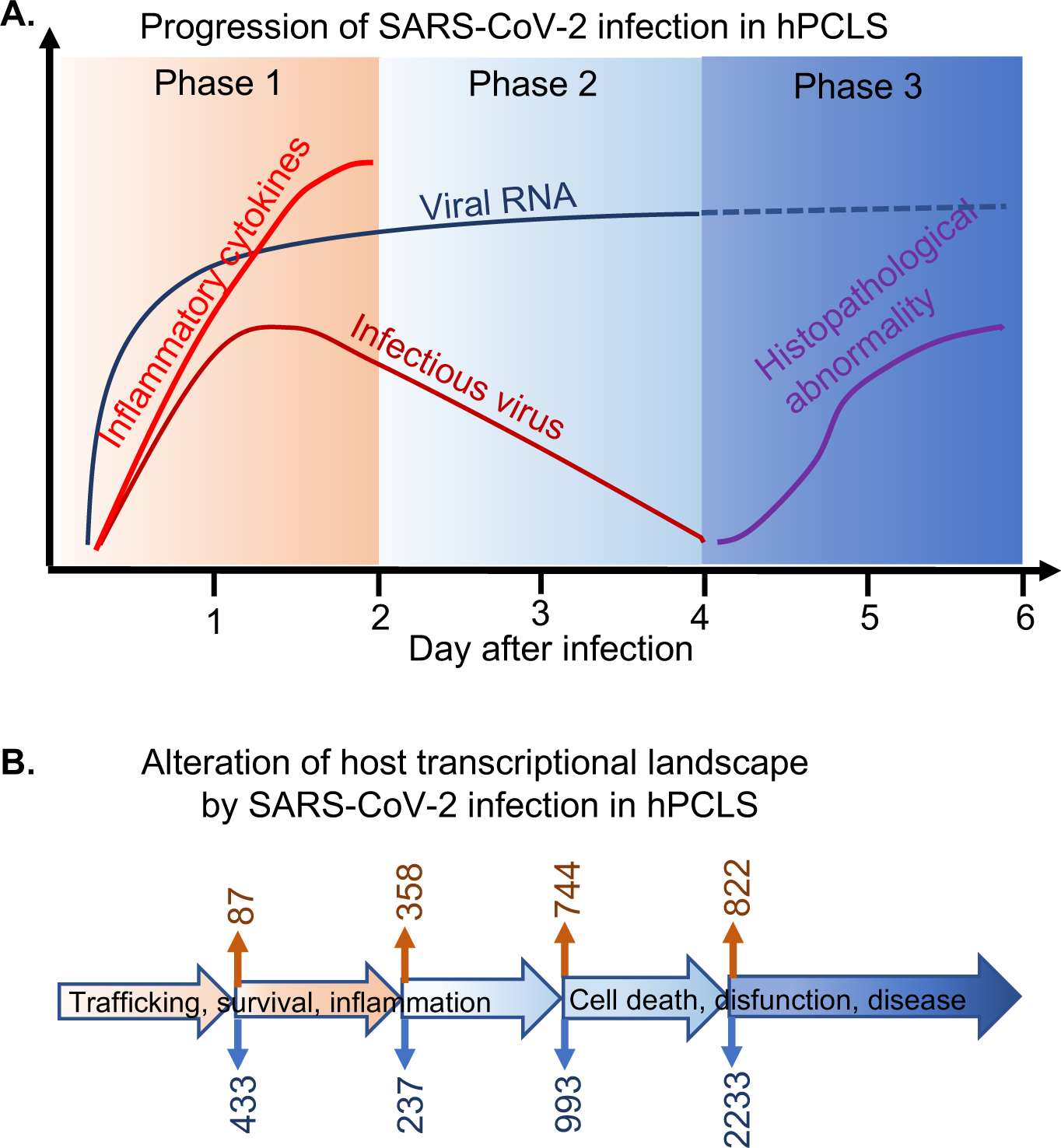
Schematic presentation of the pathogenic progression following SARS-CoV-2 infection in hPCLS. (A) The pathogenic progression is divided into 3 phases. Phase 1 is the initiation phase, represented by rapid virus replication and induction of proinflammatory cytokines and chemokines. Phase 2 highlights the inverse correlation between viral RNA replication and infectious virus production with no obvious macrostructural changes in the lungs. Phase 3 illustrates the consequence of SARS-CoV-2 infection in the lungs, as characterized by the development of histopathogical abnormality. Dashed line for viral RNA in this phase is the prediction. (B) Alteration of the host transcriptional landscape by SARS-CoV-2 infection. The number of genes that are up-and down-regulated by SARS-CoV-2 infection at indicated timepoints as in (A) are shown at the top (orange) and bottom (blue), respectively. The overall transcriptional landscape appears switching from promoting trafficking, survival and inflammation at the initial phase to cell death, disfunction, and disease in the second phase, which may drive the disease progression to the final phase.

IFN-I are key antiviral cytokines that can be induced upon infection by diverse viruses and play an important role both in host defense and in mediating inflammation. Hence fine tuning of the IFN-I responses often dictates the outcome of the infection or disease, and dysregulation of the IFN-I signaling pathways often contributes to disease pathogenesis. In our transcriptomic profiling, we found a complete absence of, or even negative, IFN-I response to SARS-CoV-2 infection in hPCLS (**Fig. 5G, Dataset S1-S4**), suggesting a protective role of IFN-I in COVID-19 pathogenesis. This interpretation is supported by the identification of impaired IFN-I responses in severe COVID-19 patients that preceded clinical worsening (39), and genetic mutations in Toll-like receptor (TLR)-3-dependent and IRF7-dependent IFN-I immunity (40), TLR7 deficiency (41), or autoantibodies against IFNα, IFNβ, and IFNω (42), as major risk factors for the development of severe COVID-19 (43, 44). Defective activation and regulation of IFN-I immunity has also been linked to increased COVID-19 severity as evidenced in postmortem lung tissues from lethal cases of COVID-19 (45) and in peripheral blood of patients with severe COVID-19 (39, 46). Further, upregulated IFN-I responses in asymptomatic COVID-19 infection are associated with improved clinical outcome (47). However, robust IFN-I responses have also been reported in peripheral blood mononuclear cells (PBMCs) from patients with severe COVID-19 (48–50), in bronchoalveolar lavage fluid from COVID-19 patients (51), and in human lung stem cell-based alveolospheres following SARS-CoV-2 infection (13). The apparent contradictory roles of IFN-I responses during SARS-CoV-2 infection might be explained by a number of variables, such as the type of cells and tissues being analyzed, the methods being used, the timing of the sample collection, and specific subsets of IFN-I or interferon-stimulated genes. Our transcriptomic profiling revealed a striking similarity in IFN-I signaling in hPCLS after SARS-CoV-2 infection and in postmortem lung tissues from lethal COVID-19 patients (45). Therefore, the hPCLS may provide an ideal platform for further delineating the roles and mechanisms of IFN-I responses in COVID-19 pathogenesis.

Using the hPCLS platform we confirmed the antiviral activity of HHT against SARS-CoV-2 and identified parameters that can help evaluate its therapeutic effect on the clinical outcome of COVID-19, as evidenced by the inhibition of inflammatory cytokine and chemokine expression and the elimination of histopathological abnormalities caused by SARS-CoV-2 infection (**Fig. 7**). Although it is not known whether HHT inhibits the induction of these cytokines directly or through inhibition of SARS-CoV-2 replication indirectly, the induction of IL-1β following virus infection and HHT treatment compared to virus infection alone suggests that HHT may have a direct effect on cytokine induction. This assumption is further supported by the evidence that HHT reduced the level of IP-10 in mock-infected hPCLS or reduced IP-10 in SARS-CoV-2-infected hPCLS to a level even lower than in mock-infected and mock-treated hPCLS (**Fig. 7B**). The varying degrees of innate immune responses among different donor hPCLS to SARS-CoV-2 infection and HHT treatment likely reflect human population heterogeneity. Thus, the hPCLS platform has many advantages over traditional cell culture systems for preclinical testing of antiviral drugs, as the hPCLS platform can evaluate antiviral efficacy as well as host factors involved in pathogenesis and potential side-effects of a given drug. It is worth noting that some of the hPCLS used in this study are revived from cryopreserved lungs, demonstrating the feasibility of using cryopreserved tissues. This allows the establishment of a “library” of donated lungs for continuous antiviral drug testing, which would resemble a clinical trial at the bench.

In summary, we have established the utility of hPCLS as an infection platform for studying SARS-CoV-2 pathogenesis and for evaluating the efficacy of antiviral drugs. We showed that during the initial stage of infection SARS-CoV-2 replicates rapidly in hPCLS, concomitant with a rapid induction of multiple pro-inflammatory cytokines and chemokines, which is consistent with the observations from COVID-19 patients. At the late stage, infectious viruses decreased rapidly while viral RNAs persisted and histopathological changes ensued. Transcriptomic and proteomic analyses identified molecular signatures and cellular pathways that are largely consistent with the disease progression. Furthermore, we have demonstrated that HHT is an effective antiviral that limits SARS-CoV-2 replication, may modulate host inflammatory responses to the advantage of the host and ameliorates histopathological abnormality caused by SARS-CoV-2 infection.

## MATERIALS AND METHODS

### Virus and cells

SARS-CoV-2 strain USA-WA1/2020 was obtained through BEI Resources, NIAID, NIH, and was propagated in Vero cells. A549-hACE2 cells were provided by Ralph Baric (University of North Carolina at Chapel Hill). All cells were cultured in DMEM (Gibco) containing 10% fetal bovine serum (FBS) and 1% penicillin and streptomycin at 37 °C in 5% CO_2_.

### Preparation of human precision-cut lung slices (hPCLS)

Lungs were obtained from anonymous donors through transplant teams of the Arkansas Regional Organ Recovery Agency (ARORA) and by the National Disease Research Interchange (NDRI). Lung vasculature was perfused with phosphate-buffered saline (PBS) to wash out residual blood. The lobes were surgically dissected, and the major bronchi were cannulated. Individual lobes were inflated with sterile 1.8% low-gelling-temperature agarose in PBS at 37°C. After inflation, bronchi were clamped and incubated at 4 to 7 °C for 2 to 3 hours to allow the agarose to solidify. Hardened lungs were cut into ∼12-mm-thick sections, and cross-sectioned airways were identified and collected with an 8.5-mm-diameter coring tool under a dissecting microscope. The cores (80–100 per lobe) were further cut into 600-µm-thick slices. Slices were then cultured in 48-well plates in DMEM/Ham’s nutrient mixture F-12 medium (DMEM-F12; 1:1) supplemented with 10% FBS, antibiotic-antimycotic, and antibiotic formulation (Primocin) at 37 °C in 5% CO_2_ with continuous agitation in a humidified incubator. hPCLS were cultured for 4-5 days before they were used for infection.

### Infection of hPCLS and determination of virus titer

hPCLS were infected with SARS-CoV-2 at 2×10^5^ TCID_50_ per slice in a 48-well tissue culture plate at 37 °C. At 1 h p.i., the virus inoculums were removed and hPCLS were rinsed with 1x PBS. hPCLS were cultured for various periods of time as indicated. Virus titers in the supernatants were determined by the standard TCID_50_ assay on Vero cells, and were expressed as TCID_50_/ml.

### Immunofluorescence assay and hematoxylin-eosin staining of hPCLS

hPCLS were fixed with 10% formalin for 30 min at room temperature, and then processed, embedded with paraffin, sectioned and stained with hematoxylin-eosin (H&E) at the Experimental Pathology Core facility at UAMS. Unstained slides were used for detection of viral proteins by immunofluorescence assay using the primary monoclonal antibody against SARS-CoV-2 nucleocapsid (N) protein (1:50 dilution) (BEI Resources, NR-619) and the secondary anti-mouse IgG antibody conjugated with FITC (1:100 dilution, Sigma). The cell nuclei were stained with DAPI. The slides were observed under a fluorescence microscope (Olympus IX-70), and images were captured with the attached digital camera (Zeiss).

### Cytokine measurements

hPCLS culture supernatants were inactivated for SARS-CoV-2 prior to cytokine assay. Cytokines were measured by flow cytometry using cytometric bead array (CBA) kits (BD Biosciences) following the manufacturer’s instruction. The human inflammatory cytokine kit (IL-8, IL-1β, IL-6, IL-10, TNF-α, IL-12p70) (Cat.# 551811) and human chemokine kit (CXCL8/IL-8, CXCL9/MIG, CXCL10/IP-10, CCL2/MCP-1, CCL5/RANTES) (Cat.# 552990) were used. The amount of each cytokine/chemokine in virus-infected hPCLS was normalized to that in mock-infected hPCLS and was expressed as pg/ml.

### RNA isolation and quantitative reverse transcription and polymerase chain reaction (qRT-PCR)

Total RNAs were isolated from hPCLS using TRIzol reagent (Invitrogen) according to the manufacturer’s instruction. qRT-PCR was carried out using iScript RT Supermix (BioRad cDNA kit, cart#1708841) and iTaq Universal SYBY green Supermix kit (BioRad cat# 1725121) in a thermal cycler (QuantStudio 6 Flex, Applied BioSystems) according to the manufacturer’s instruction (BioRad). The primer pair specific to SARS-CoV-2 N gene (forward primer: 5’-ATG CTG CAA TCG TGC TAC AA-3’; reverse primer: 5’-GAC TGC CGC CTC TGC TC-3’) or to cellular housekeeping gene GAPDH (forward primer: 5’ TGA TGA CAT CAA GAA GGT GGT GAA G −3’; reverse primer: 5’TCC TTG GAG GCC ATG TGG −3’) were used for amplifying viral and cellular RNA, respectively. The amount of viral RNA was normalized to that of GAPDH and expressed as fold change relative to mock-infected sample.

### Gene expression profiling by RNAseq

The RNA samples isolated from hPCLS were sequenced by Novogene (www.novogene.com). The reads were mapped using STAR (v2.5) (52) to the reference genome and HTSeq (v0.6.1) (53) and used to count the reads mapped to each gene. FPKM of each gene was calculated and differential expression analysis performed using DESeq2 (v2_1.6.3) (54). The resulting p-values were adjusted using the Benjamini and Hochberg’s approach for controlling the False Discovery Rate (FDR). Genes with an adjusted p-value <0.05 were considered differentially expressed. Gene Ontology (GO) enrichment analysis of differentially expressed genes was performed using clusterProfiler R package to test the statistical enrichment of differential expression genes in KEGG pathways (55, 56). Volcano plots were created using VolcaNoseR (57).

### Proteomic analysis

Proteins were isolated from hPCLS with the radioimmunoprecipitation assay (RIPA) buffer (Thermo, Cat.# 89901) containing a cocktail of protease inhibitors (Sigma, Cat.# P8340-5ML) and phosphatase inhibitors (Fisher, Cat.# PIA32957) at room temperature for 30 min followed by repeated pipetting. Proteins were analyzed by using CME bHPLC phosphoTMT Methods – Orbitrap Eclipse in the IDeA National Resource for Quantitative Proteomics facility on UAMS campus (58–61). Proteins and phosphopeptides with an FDR-adjusted p-value <0.05 and an absolute fold change >2 were considered significant.

### Antiviral drug testing

hPCLS were infected with SARS-CoV-2 at 2×10^5^ TCID_50_ per slice and treated with homoharringtonine (HHT) (Sigma, cat# SML-1091-10MG) at 1 h p.i. at various concentrations. hPCLS treated with vehicle (medium containing 0.1% or 0.01% DMSO) were used as a negative control. At 48 h p.i., culture supernatants were collected for determination of virus titer by TCID_50_ assay and for measuring the cytokines and chemokines using the human inflammatory cytokine and chemokine CBA kits as described above.

### Cell viability assay

Cell viability was determined using the XTT assay kit TOX2-1KT according to the manufacturer’s instruction (Sigma). Medium containing DMSO at 1% or less was used as vehicle control.

### Statistical analysis

Statistical analyses on cytokine data were performed using the Tukey’s multiple comparisons test in the GraphPad Prism 9 program (v9.5.0). Other statistical analyses were carried out with unpaired T test or one way ANOVA in the same program. Results with *P* values of >0.05, <0.05, <0.01, <0.001, and < 0.0001 are indicated in the figures and legends.

### Data Availability

All data supporting the findings of this study are found within the paper and its Supplemental Figures and Datasets, and are available from the corresponding author upon request. RNAseq data have been deposited in the Gene Expression Omnibus (GEO) database under accession GSE226702. The proteomics data have been deposited in the MassIVE repository with accession link ftp://massive.ucsd.edu/MSV000091383/.

## ACKNOWLEDGEMENT

We thank Suzanne House, Claire Putt, and Dana Frederick in the Cell Biology Laboratory, Arkansas Children’s Research Institute for processing the hPCLS. This work was supported by a seed grant from the Vice Chancellor for Research and Innovation. IDeA National Resource for Quantitative Proteomics is supported by NIH/NIGMS grant R24GM137786. The UAMS Bioinformatics Core is supported by the Winthrop P. Rockefeller Cancer Institute and NIH/NIGMS grant P20GM121293. SDB is supported by National Science Foundation Award No. OIA-1946391.

## AUTHOR CONTRIBUTIONS

Concenptualization: RDP, RCK, XZ Experimental design: RDP, XZ Methodology: RDP, PAM, SKB, RCK, XZ Investigation: RDP, PAM, SKB, SDB, DHA, AG, RCK, JLK, XZ Supervision: XZ Writing-original draft: XZ Writing-review & editing: RDP, PAM, SKB, SB, DHA, AG, RCK, JLK, XZ

